# Neuronal or vascular receptive fields? On the relation between BOLD-fMRI population receptive field (pRF) estimates and cortical vascularization

**DOI:** 10.1101/2025.06.03.657640

**Authors:** W. Schellekens, M. Nota, I. Groen, G. Piantoni, J. Winawer, N. Petridou

## Abstract

This study investigates the contribution of different vascular compartments on population receptive field (pRF) size estimates within the early visual cortex (V1, V2, and V3) using BOLD-fMRI. We employed T2*-weighted gradient-echo (GE) and T2-weighted spin-echo (SE) sequences at 7 Tesla (7T) and a multi-band GE sequence at 3T to explore how different vessel sensitivities across these sequences influence pRF modeling. Our results confirm the expected pRF size increase across eccentricity and visual areas but found no significant differences in pRF size estimates across MR sequences, voxel sizes, or field strengths. BOLD signal amplitudes were influenced by MR sequence, with largest signal changes observed for 7TGE, and amplitude increases were noted across cortical depth for GE sequences but not for SE sequences. Contrary to our hypotheses, pRF size estimates were not noticeably affected by the local vascularization, suggesting that pRFs primarily reflect neuronal activity rather than vascular compartment characteristics. Our study highlights the robust nature of pRF size estimates across various fMRI conditions and points toward the decoupling of pRF properties from vascular factors.

## Introduction

Functional magnetic resonance imaging (fMRI) is the most commonly used non-invasive technique to measure and localize human brain activity (Bandettini, 2012; Ogawa et al., 1990). Despite its widespread implementation in both scientific and clinical settings, the signal obtained from fMRI is actually a proxy for the underlying neuronal activity. In most fMRI studies, the signal relies on the blood-oxygenation-level-dependent (BOLD) contrast, which capitalizes on changes in the ratio of paramagnetic deoxyhemoglobin to diamagnetic oxyhemoglobin in venous blood, reflective of changes in metabolic demands in relation to neuronal activity (Buxton, 2012; Buxton et al., 2004; Gauthier and Fan, 2019; Havlicek and Uludağ, 2020; Logothetis and Pfeuffer, 2004). Because the amplitude of the BOLD signal scales with increased oxygenated blood volume, BOLD signal properties in general depend on the total blood volume, which in turn depends on local vessel properties such as vessel size and reactivity (Gati et al., 1997; Roefs et al., 2024; Schellekens et al., 2023; Siero et al., 2013; Uludaǧ et al., 2009). However, it is not well-defined how signals originating from different cortical vascular compartments affect inferences on brain function. In the current study, we investigate the effect of relatively larger (here referred to as macro vessels containing pial and intracortical veins) and smaller vascular compartments (referred to as micro vessels consisting of capillaries, venules and arterioles) on neural metrics for studying brain function.

To investigate the interplay between vascular compartment sizes and neural metrics for studying brain function using BOLD fMRI, we focus on receptive field measurements of neurons within the early visual cortex, specifically V1, V2, and V3. Through electrophysiological recordings, neurons in the early visual cortex are known to exhibit receptive fields: the visual space that elicits a neuronal response (Gilbert and Wiesel, 1992; Hubel and Wiesel, 1959, 1977; Victor et al., 1994a). Notably, a neuron’s receptive field center corresponds to its preferred visual field location, as indicated by response amplitude. The size of a neuron’s receptive field inversely relates to its specificity in processing input. Receptive field modeling has revealed fundamental features of mammalian visual cortical architecture. Particularly, the distribution of receptive field centers has revealed multiple topographic representations of the retina (i.e., retinotopy) (Fox et al., 1987; Sereno et al., 1995; Shipp et al., 1995; Woldorff et al., 1997). Additionally, receptive field size distributions have revealed functional and hierarchical properties of visual information processing: smallest receptive field sizes have been reported in granular layer IV, which receives information directly from the thalamus (Fracasso et al., 2016; Gilbert and Wiesel, 1992; Martinez et al., 2005; Self et al., 2019); the gradual receptive field size increase from foveal to peripheral visual field locations indicates the designated processing specificity (A T Smith et al., 2001; Wilson and Sherman, 1976); and the gradual receptive field size increase from V1 to V2, V3, etc. signifies the cortical hierarchy of visual information processing (Felleman and Van Essen, 1991; A. T. Smith et al., 2001; Van Essen et al., 1992; Wandell and Winawer, 2011; Wandell et al., 2015). Receptive field analyses have successfully been deployed in fMRI studies, measuring the aggregate receptive field of a population of neurons within a single voxel: population Receptive Field (pRF) modeling (Dumoulin and Wandell, 2008; Fracasso et al., 2016; Wandell and Winawer, 2015; Zuiderbaan et al., 2012). Thus, receptive field modeling has been successfully deployed using various measuring modalities and, therefore, provides an optimal metric to investigate MR-dependent effects of local vascularization on neural inferences.

In this study, we employ both gradient-echo (GE) and spin-echo (SE) BOLD-fMRI at 7T, as well as GE-BOLD-fMRI at 3T, to quantify the impact of vascular compartment sizes on pRF estimates, including the influence of intra- and extra-vascular components. The spatial specificity of these imaging techniques depends on their sensitivity to different vessels; the greater an imaging sequence’s sensitivity to larger vessels, the more it reflects blood outflow from spatially extended regions, thereby lowering its effective spatial resolution (Turner, 2002). Among these techniques, the T2-weighted SE BOLD-fMRI contrast supposedly offers greatest spatial specificity (BOLD point-spread function (PSF) < 2 mm full-width at half-maximum (FWHM), (C. A. Olman et al., 2004; Laura M. Parkes et al., 2005), although precise SE-BOLD PSF estimates at 7T do currently not exist. SE-BOLD’s high spatial specificity is due to the 180-degree refocusing pulse of the SE BOLD-fMRI sequence, which largely nullifies large physiological contributions (e.g., T2* effects from large veins), leaving a microvascular-weighted BOLD effect (i.e. primarily sensitive to smaller vessels) that is relatively evenly distributed across cortical depth and thought to closely correspond to the neuronal source (Markuerkiaga et al., 2016; Norris, 2012; Norris et al., 2002; Oja et al., 1999; Roefs et al., 2024; Schellekens et al., 2023; Yacoub et al., 2003). In contrast, the GE-BOLD signal is sensitive to all venous vascular compartments (capillaries, venules, and large draining veins) and scales with vessel diameter, meaning larger vessels exhibit larger signal changes compared to smaller vessels (Siero et al., 2013; Uludaǧ et al., 2009; Zhao et al., 2004). As a result, GE-BOLD has a lower spatial specificity than SE-BOLD as represented by a larger PSF. Previous PSF estimates for GE-BOLD vary between 0.8 and 1.8mm (Fracasso et al., 2021) and approximately 2mm (C A Olman et al., 2004; Shmuel et al., 2007) at high field strengths (≥7 Tesla) and are substantially larger at 3 Tesla (FWHM = 3.9 mm) (Laura M Parkes et al., 2005). This difference in GE-BOLD PSF between higher and lower field strengths is primarily due to the reduction of intravascular effects at high field strengths, where the T2 of blood is too short to impact most BOLD-fMRI measurements. Consequently, BOLD-fMRI contrast at 7T primarily reflects extra-vascular effects, which theoretically align more closely with underlying neuronal function. In contrast, BOLD-fMRI measurements at 3T or lower field strengths include intravascular effects, further decreasing spatial resolution and the accuracy of neuronal activity assessment (Jochimsen et al., 2004).

In exploring the relationship between vascular compartment sizes and neural metrics (such as pRF estimation) as measured by BOLD fMRI, we propose several hypotheses. We anticipate that the estimated pRF sizes will vary depending on the vascular compartments contributing to the estimate, even though the underlying neuronal receptive field properties remain constant across imaging sequences. Generally, we expect microvascular compartments to produce smaller receptive field sizes compared to macrovascular compartments. Although a recent imaging study reported similar pRF sizes between the Vascular-Space-Occupancy (VASO) sequence—believed to be more sensitive to microvascular compartments—and 7T GE-BOLD (Oliveira et al., 2022), we hypothesize that pRF size estimates will differ due to the reported spatial specificity differences in deoxyhemoglobin-based BOLD fMRI sequences. Additionally, we hypothesize that there will be differences in pRF sizes across cortical depth when comparing 7TSE and 7TGE, given their distinct sensitivities to local vascular compartments.

## Methods

### Participants

Ten healthy volunteers (N = 10, age range 20-26 years, M = 22.9, SD = 2.07, five females) participated in the study after giving written informed consent. All participants had normal or corrected-to-normal visual acuity. The experimental protocol was approved by the ethics committee of the University Medical Center Utrecht (UMCU), and conducted conforming to the principles of the 2013 Declaration of Helsinki and in accordance with the Dutch Medical Research Involving Human Subjects Act.

### Visual field map stimulus

The visual field map stimulus consisted of sweeping bar apertures in eight directions (4 cardinal and 4 diagonal directions), exposing a static greyscale bandpass noise pattern (Zhou et al., 2018). The stimuli were windowed within a circular aperture and positioned at the center of the visible part of the screen; the rest of the display was filled with neutral grey. Individual sweeping bar apertures were presented for 500ms on screen and followed by a 350ms inter-stimulus interval, after which the next location was presented on screen or a blank period of neutral grey ensued. During 1 mapping run, 8 sweeps across the visual field were performed in 212.5s, albeit that the first and last 11.9s were blank periods by default. During each scan session, 2 visual field mapping experiments were conducted. The stimuli were generated in MATLAB (Mathworks, Natick, MA, USA) using PsychToolbox. At the 3T scanner, the stimuli were displayed on a 32.4 × 51.8 cm LCD monitor (1920×1200 pixels at 60 Hz), located at the head of the scanner bed at a distance of approximately 142 cm from the subjects’ eyes, and were viewed through an angled mirror mounted on the head coil. The maximal radius of the circular stimulus was 4.42° visual angle, occupying the height of the display. At the 7T scanner, the stimuli were back-projected onto a 13.0 × 8.0 cm screen (1920×1080pixels at 60 Hz) at a viewing distance of approximately 38 cm and were observed through prism glasses and a mirror. Here, the radius of the stimulus was 4.42° visual angle as well. In order to keep the subjects vigilant they were tasked with fixating on a cross (0.03° visual angle), positioned at the center of the display, and reporting its change in color by pressing an MR-compatible button box. The color alternated between red and green at random intervals unrelated to the stimulus sequence.

### MRI protocols

Scanning was performed at the UMCU, Utrecht, The Netherlands, on a 3T Philips Ingenia scanner with a 32-channel receive head coil (Philips Healthcare, Best, The Netherlands) and a 7T Philips Achieva scanner with two 16-channel high-density surface receive arrays (MRCoils BV, The Netherlands) (Petridou et al., 2013). All subjects completed three separate scanning sessions: one at 3T and two at 7T.

Whole-brain anatomical (T1-weighted) data were collected at 3T at 1.0 mm^3^ isotropic voxel size, field-of-view (FOV) (anterior-posterior (AP), foot-head (FH), right-left (RL)) = 232 × 256 ×192 mm, repetition time (TR) = 7.9ms, echo time (TE) =4.5ms. At both 7T scanning sessions, T1-weighted MPRAGE volume was recorded with the following specifications: FOV (AP, FH, RL) = 40 x 159 x 159 mm, which covered the posterior part of the brain (occipital lobe/early visual cortex), voxel size: of 0.8 x 0.8 x 0.8 mm, and TR/TE = 7.0/2.97 ms. T1-weighted volumes at high field strength can experience substantial intensity inhomogeneities. Therefore, a proton density (PD) volume of equal dimensions was acquired to correct for these large-scale intensity inhomogeneities.

All BOLD fMRI measurements were acquired using an echo-planar imaging (EPI) sequence. The functional scans at 3T were obtained using a gradient-echo (GE) EPI multiband sequence with the following parameters: multiband factor = 2, SENSE factor = 3.0, TR/TE = 850/35ms, flip angle (FA) = 60°, number of slices = 30, FOV (AP, FH, RL) = 60 × 192 ×192 mm, voxel size = 2.0 mm isotropic. At 7T, the functional data of the first session were collected using a GE-EPI sequence with SENSE factor =3.0, TR/TE = 850/27ms, FA = 60°, number of slices = 7, FOV (AP, FH, RL) = 7 × 128 × 128 mm covering parts of early visual cortex, voxel size = 1.0 mm isotropic. The functional data of the second 7T session were acquired with a SE-EPI sequence with the following settings: SENSE factor = 2.0, TR/TE= 850/50ms, FA = 90°, number of slices = 5, FOV (AP, FH, RL) = 7.5 ×190 × 190 mm, voxel size = 1.5 mm isotropic. For each session, an additional 10 volumes were acquired with the phase encoding direction reversed, but otherwise identical parameter settings. During each fMRI session, two visual field mapping runs were completed, consisting of 250 volumes each (duration = 212.5s), resulting in 500 volumes per fMRI session, per participant. In addition to the magnitude data, also the phase data from each session was reconstructed.

### Preprocessing

The 3T T1-weighted volume was used to determine regions of interest (ROI). White matter and pial surfaces were estimated using Freesurfer (https://surfer.nmr.mgh.harvard.edu), and ROIs of visual cortex were estimated on the cortical surfaces using Neuropythy (https://github.com/noahbenson/neuropythy). ROIs were projected to volumetric space using Freesurfer’s mri_surf2vol command. The volumetric ROI data were transformed onto functional space (i.e., shape and orientation of the functional data) of each functional session, using (a linear combination of) affine transformation matrices and a nearest-neighbor interpolation type.

The 7T T1-weighted volumes were divided by their corresponding PD volumes to correct for large-scale intensity inhomogeneities (Van de Moortele et al., 2009). Afterward, the T1-weighted volume was resampled to a resolution of 0.2mm3 isotropic voxel size to estimate cortical layers at high spatial resolution. Grey matter tissue was estimated using ANTs’ AtroposN4 command (Avants et al., 2011), using in-house Python scripts. Next, estimated cortical tissue was divided into 18 equivolumetric layers using the LayNii software package (Huber et al., 2021). The term ‘layers’ should not be taken to represent architectonic layers distinguishable from histology. Here, ‘layers’ merely reflect discrete cortical depth levels. The layer mask was transformed onto the functional space of the corresponding 7T session, using the appropriate affine transformation matrix and a nearest-neighbor interpolation type.

All functional time series were denoised using NORDIC on the basis of the magnitude and phase data (Moeller et al., 2021). Next, the time series were corrected for rigid body head motion with AFNI’s 3dvolreg (afni.nimh.nih.gov). Geometric distortions caused by EPI-dependent susceptibility effects were corrected for using AFNI’s 3dQwarp, estimating the geometrical mid-point of the mean time series and the mean of the reversed-phase time series. The susceptibility distortion correction and the motion correction were simultaneously applied in a single interpolation step using 3dNwarpApply to generate motion-corrected undistorted functional time series (Cox J.S., 1996). At this stage we, additionally resampled the 7TGE to 2.0mm isotropic voxel using ANTs’ ResampleImage with linear interpolation, matching the 3TGE voxel size, for comparison purposes. Several affine registrations were then performed between the mean volume of each functional time series and the corresponding T1-weighted volume, and between the T1-weighted volumes of each scan session, using antsRegistration (http://stnava.github.io/ANTs/). The affine transformation matrices were used to transform previously computed layer masks, and V1, V2, V3 ROI masks to the origin and dimensions of the functional volumes using nearest-neighbor interpolation. The functional time series were then high-pass filtered using a discrete cosine transform filtering with a cut-off at 0.01 Hz and rescaled to percent signal change (%BOLD). As a final step, the time series were temporally smoothed using a Savitzky-Golay filter with a window length of 11s and 4th order polynomials (Savitzky, A.; Golay, 1964; Schmid et al., 2022).

### pRF analysis

Population Receptive Field estimates were calculated using a custom Python code, computing stimulus-dependent model time series. For each voxel within the layer- and ROI-masks, model time series were generated on the basis of the dot product of the stimulus apertures over time and a gaussian function describing a population receptive field convolved with a canonical hemodynamic response function (HRF) (1).

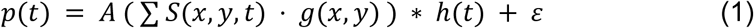

where, S(x,y,t) is the stimulus aperture across 2-dimensional visual space and time, g(x,y) is the elicited response at visual field location x,y following a Gaussian shaped receptive field (see equation 2). h(t) is a canonical HRF, A controls the signal amplitude, and ε is the model error.

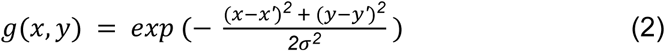

where, x’ and y’ represent the receptive field centers along the horizontal and vertical axes, respectively, and σ reflects the receptive field size. This procedure of finding pRF fits uses the Levenberg-Marquardt linear least-square minimization algorithm, and produces two pRF centers (x’ and y’), a pRF size, and a mean time series amplitude per voxel. The Cartesian receptive field centers were transformed to polar coordinates (i.e. polar angle and eccentricity). The full model fit is also used to calculate *R-squared (r^2^)* values.

### pRF models

We developed population receptive field (pRF) models for various scan sequences at their respective magnetic field strengths, informed by previously reported point spread functions (PSFs) described as Gaussian spreads in millimeters FWHM. First, we modeled eccentricity representation as a function of cortical surface distance from the 10° visual angle mark, following (Engel et al., 1997):

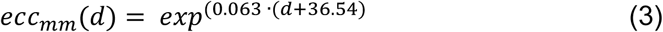

where *d* is the distance in mm on the cortical surface. Next, we calculated pRF size increases with increasing eccentricity representation in mm cortical surface (Dumoulin and Wandell, 2008; Hubel and Wiesel, 1962; Victor et al., 1994b):

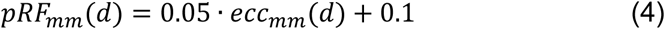

To model the spatial resolution of the scan sequences, we used the following Gaussian function:

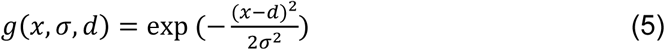

where *x* is the center of the Gaussian function, *σ* is the Gaussian standard deviation, and *d* is the distance from the center in cortical mm.

The PSFs of the scan sequences were integrated by converting FWHM values reported in the literature into Gaussian standard deviations:

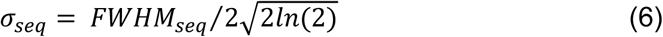

The FWHM values used were: 7TSE = 0.9mm (linear increase from WM surface 0.8mm to pial surface 0.97mm), 7TGE = 1.3mm (linear increase from WM surface 0.8mm to pial surface 1.8mm), and 3TGE = 3.9mm (Fracasso et al., 2021; C. A. Olman et al., 2004; Laura M. Parkes et al., 2005; Schiller et al., 1976). Since 7TSE PSF values are less documented, they were estimated as similar to the lowest 7TGE PSF values and assumed nearly constant across cortical depth. Using these values, we estimated pRF size as follows:

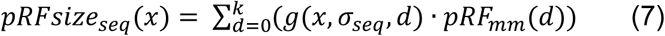

where *x* and *d* are eccentricity representations in cortical mm, with *x* reflecting preferred eccentricity and d accounting for all included representations.

This approach generated three pRF size estimates (for 7TSE, 7TGE, and 3TGE) per millimeter of cortical surface. These were then interpolated for different voxel sizes: 1.5mm, 1.0mm, and 2.0mm. Additionally, 7TGE estimates were interpolated at 2.0mm voxel size to directly compare with 3TGE.

Finally, the method was extended to create vascular compartment-dependent pRF size estimates across cortical depth. This included a small linear PSF increase for 7TSE (from 0.8mm at the white matter surface to 0.97mm at the pial surface) and a larger PSF increase for 7TGE (from 0.8mm to 1.3mm). Using these values, we estimated pRF sizes across the cortical depth of approximately 2.25mm, matching the cortical thickness of the early visual cortex (Alvarez et al., 2019; Fischl and Dale, 2000).

The resulting pRF models reflect the spatial specificity of each scan sequence, offering quantitative expectations for actual pRF measurements (Figure 1). The smallest pRF sizes are predicted for 7TSE BOLD (0.13°–0.37° visual angle across all cortical eccentricity representations, with a maximum of 5.0°), followed by 7TGE BOLD (0.18°–0.49°), and the largest for 3TGE BOLD (0.52°–1.46°).

**Figure 1.**
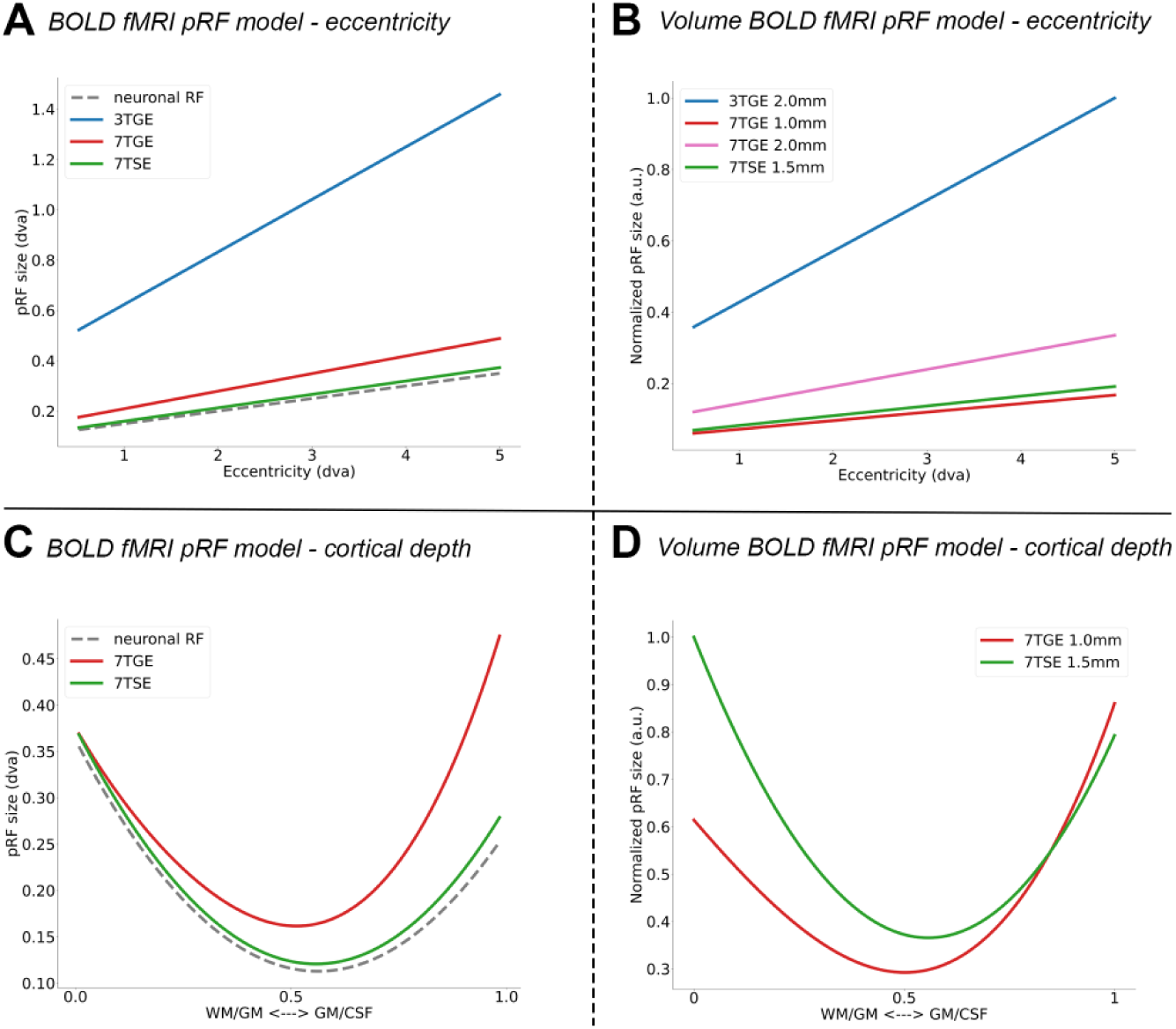
pRF models. A) pRF size increase over eccentricity per MR scan sequence. The point-spread function of each MR-sequence determines the deviation from the assumed neuronal receptive field size. B) same as A) with the applied voxel sizes taken into account. Additionally, a pRF model was made for the 7TGE acquisition with increased voxel size (2.0mm; pink line). The pRF sizes were normalized with respect to the maximum modeled pRF size. C) pRF size variation across cortical depth for the 7T scan sequences. The point-spread function increases linearly across cortical depth (from WM/GM to GM/CSF) for 7GE and remains nearly constant for 7TSE. The appropriate voxel sizes are taken into account in D).

When voxel size is considered, pRF sizes for 1.5mm isotropic voxels at 7TSE are about 13% larger than those for 1.0mm isotropic voxels at 7TGE. This suggests that differences in voxel size influence observed pRF size differences between 7TGE and 7TSE. However, the absolute differences between these modalities remain small. Larger differences are expected between 7T and 3T, where 3TGE pRF sizes are approximately 5–6 times larger than 7TGE and 7TSE at 2.0mm isotropic voxel size. Even when 7TGE voxel size is increased to 2.0mm, 3TGE pRF sizes remain roughly 3 times larger. These differences reflect the influence of intravascular effects on pRF size estimation (Figure 1B).

Across cortical depth, the models predict that 7TGE pRF sizes exceed 7TSE, with differences of up to 70% near the pial surface, where 7TGE pRF estimates are affected by large pial veins (Figure 1C). Conversely, in deeper layers near the white matter surface, 7TSE pRF sizes are predicted to be up to 60% larger than 7TGE when accounting for voxel size (Figure 1D).

### Statistical analyses

For the statistical analysis, all voxels were included for which the pRF model fit explained at least 5% of the variance (*r^2^* >= 0.05). An additional inclusion constraint was applied to voxels of the 3T MR sequence, ensuring that the included voxels spatially overlapped the FOV of the 7T MR sequences. The model parameters that were estimated for each voxel (i.e., polar angle, eccentricity, pRF size and amplitude) were not binned, but were directly added to several linear mixed effect (LME) models (Statsmodels v0.14.0). Primarily, we created two LME models; one with pRF as dependent variable and another with the BOLD amplitude (percent signal change) as dependent variable. Both LME models contained MR sequence (i.e. 7TSE, 7TGE, and 3TGE) and spatial localization in the form of visual area (i.e. V1, V2 and V3) and eccentricity representation (continuous scale ranging from 0° to 5° visual angle), including the interactions as fixed effects. Random effects were estimated for the participants. Two additional LME models were created with the cortical depth level (i.e., deep layers, middle layers, superficial layers) as an additional fixed effect for the two 7T MR sequences only. Bivariate linear regression analyses were performed for pRF size estimates among MR sequences, pRF size estimates versus BOLD amplitude estimates. Significant correlations were tested for using a one-sample t-test.

## Results

Population Receptive Field estimates were obtained from visual field map experiments, conducted at 7T using GE-EPI and SE-EPI scan sequences and at 3T using a GE-EPI scan sequence (generally referred to as MR sequences). The mean and standard deviation of the pRF model fit explained variance (*r^2^*) for 7TSE was m = 0.15, s.d. = 0.11; for 7TGE m = 0.21, s.d. = 0.14; and for 3TGE m = 0.20, s.d. = 0.15. During the visual field map experiments, participants performed an attention task with on average 80% correct button presses following a color change of the fixation dot.

### Population receptive field sizes

We found that pRF sizes significantly increased along the cortical hierarchy in included visual areas from V1 to V3 (z = 2.37, p = 0.018, β = 0.23, 95% CI = [0.04, 0.43]), and across eccentricity representations (z = 3.26, p = 0.001, β = 0.27, 95% CI = [0.11, 0.43]), which both have been documented numerous times before (Benson et al., 2018; Wandell and Winawer, 2015) (Figure 2, supplementary Figures 1-5). Importantly, pRF size estimates did not significantly differ across MR sequences (z = 0.61, p = 0.541, β = 0.07, 95% CI = [-0.16, 0.30]), despite a significant interaction effect of MR sequence, visual area and eccentricity representations on pRF size estimates (z = 2.39, p = 0.017, β = 0.04, 95% CI = [0.01, 0.08]). The latter indicates that the typical progression of pRF sizes across eccentricity representations and visual hierarchy differs depending on MR sequence (Figure3A). Lastly, we did not observe that estimated pRF sizes linearly differed across cortical depth from pial to white matter surfaces for both 7T MR-sequences (z = −1.39, p = 0.164, β = −0.07, 95% CI = [-0.17, 0.03]). However, pRF size estimates were significantly influenced by an interaction of cortical depth and the visual areas (z = 2.92, p = 0.004, β = 0.07, 95% CI = [0.02, 0.10]). The 7T MR sequences do not reveal the previously reported U-shaped pattern of pRF-sizes across cortical depth, despite depth-dependent pRF-size effects differing across cortical areas. Nor did we identify hypothesized differences at the WM/GM border and GM/CSF border based on visual inspection.

**Figure 2.**
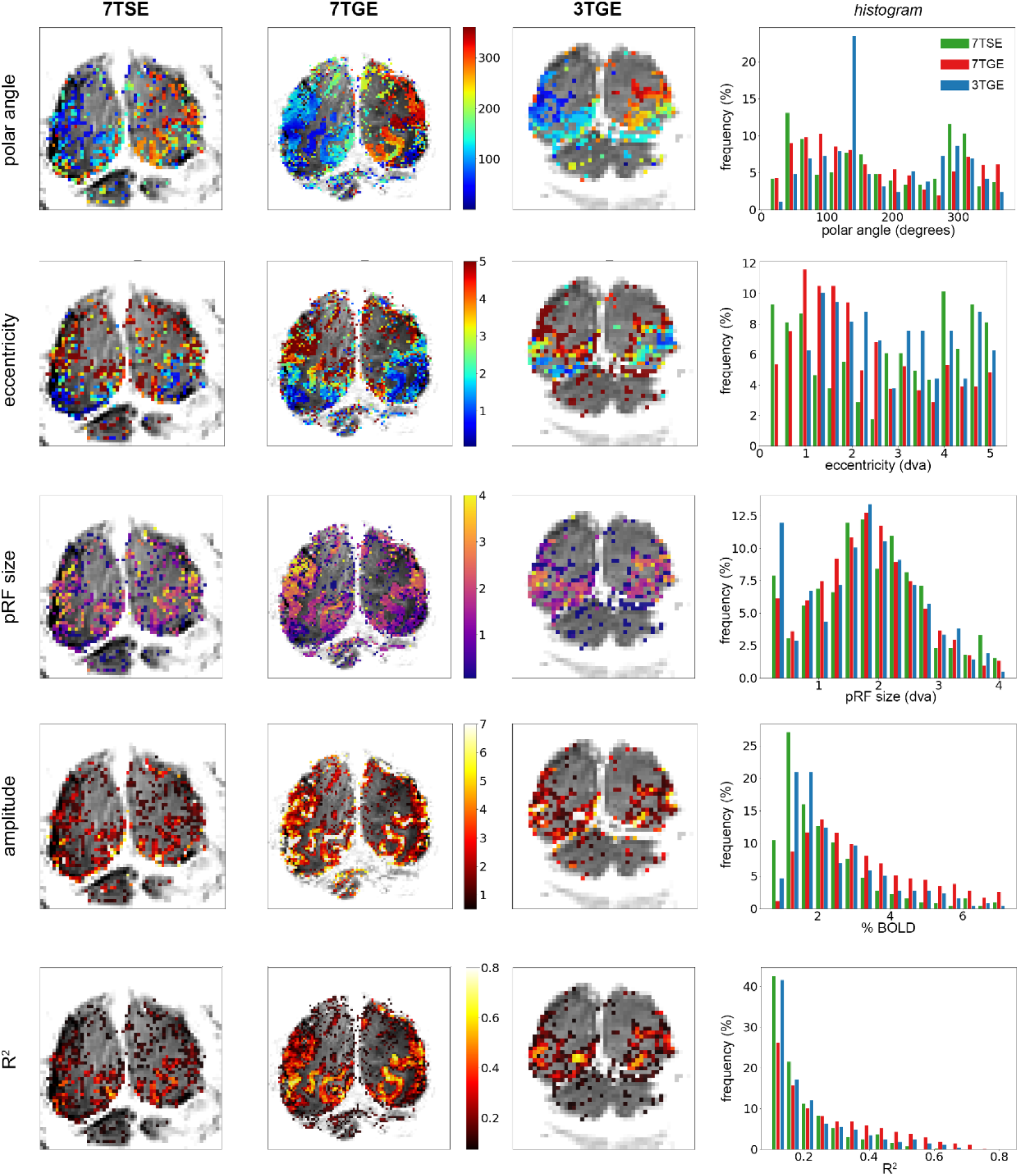
PRF fit results. pRF fitting parameters shown for each voxel in an example participant (sub-09) for each MR sequence (first three columns), including a scaled histogram of the parameter values (fourth column).

We hypothesized pRF sizes to differ across MR sequences, given each MR sequence’s unique vascular sensitivity profile. However, we did not observe a difference in pRF size estimates across MR sequences. We, additionally, would have expected a difference in pRF sizes on the basis of the different voxel sizes: larger voxels are should be sensitive to more vessels. Furthermore, the 3TGE should also be sensitive to intra-vascular effects. We expect intra-vascular effects of 3TGE scans to be apparent when comparing to 7TGE of equal voxel size. We performed the pRF model fit on the down sampled 7TGE-2.0mm scans to investigate the effect of voxel size on pRF size, and intra-vascular effects on 3TGE pRF size estimates. We did observe that pRF sizes overall increased for the 7TGE-2.0mm sequence, albeit that the increase was not significant either in comparison with the original 7TGE-1.0mm MR sequence (z = 0.46, p = 0.646, β = 0.08, 95% CI = [-0.25, 0.40]), or the 3TGE-2.0mm MR sequence (z = 0.66, p = 0.510, β = 0.30, 95% CI = [-0.59, 1.18]). These results indicate that pRF size estimates in early visual cortex are not strongly affected by voxel size for the spatial resolutions investigated here (Figure 4).

### BOLD amplitude

The BOLD percent signal change exhibited a different pattern: we did not observe a significant effect of visual areas (z = 1.15, p = 0.249, β = 0.22, 95% CI = [-0.16, 0.60]) or eccentricity representation (z = =0.89, p = 0.374, β = −0.14, 95% CI = [-0.45, 0.17]) on the estimated BOLD amplitude (Figure 3B). However, the MR sequence influenced the estimated amplitude significantly (z = 4.09, p < 0.001, β = 0.93, 95% CI = 0.49, 1.38]), reflective of relatively largest percent BOLD signal change for the 7TGE MR sequence. Lastly, we did not observe a significant effect of cortical depth level on the BOLD amplitude for the 7T MR sequences (z = −0.75, p = 0.455, β = −0.21, 95% CI = [-0.76, 0.34]), although there was a significant interaction effect of cortical depth level and MR sequence on BOLD amplitude (z = 3.35, p = 0.001, β = 0.48, 95% CI = [0.20, 0.75]). This latter result reflects the previously reported, well-documented BOLD amplitude increase from white matter toward pial surface particularly for 7TGE MR sequences but not for 7TSE MR sequences (Beckett et al., 2020; Fracasso et al., 2018; van Dijk et al., 2020).

**Figure 3.**
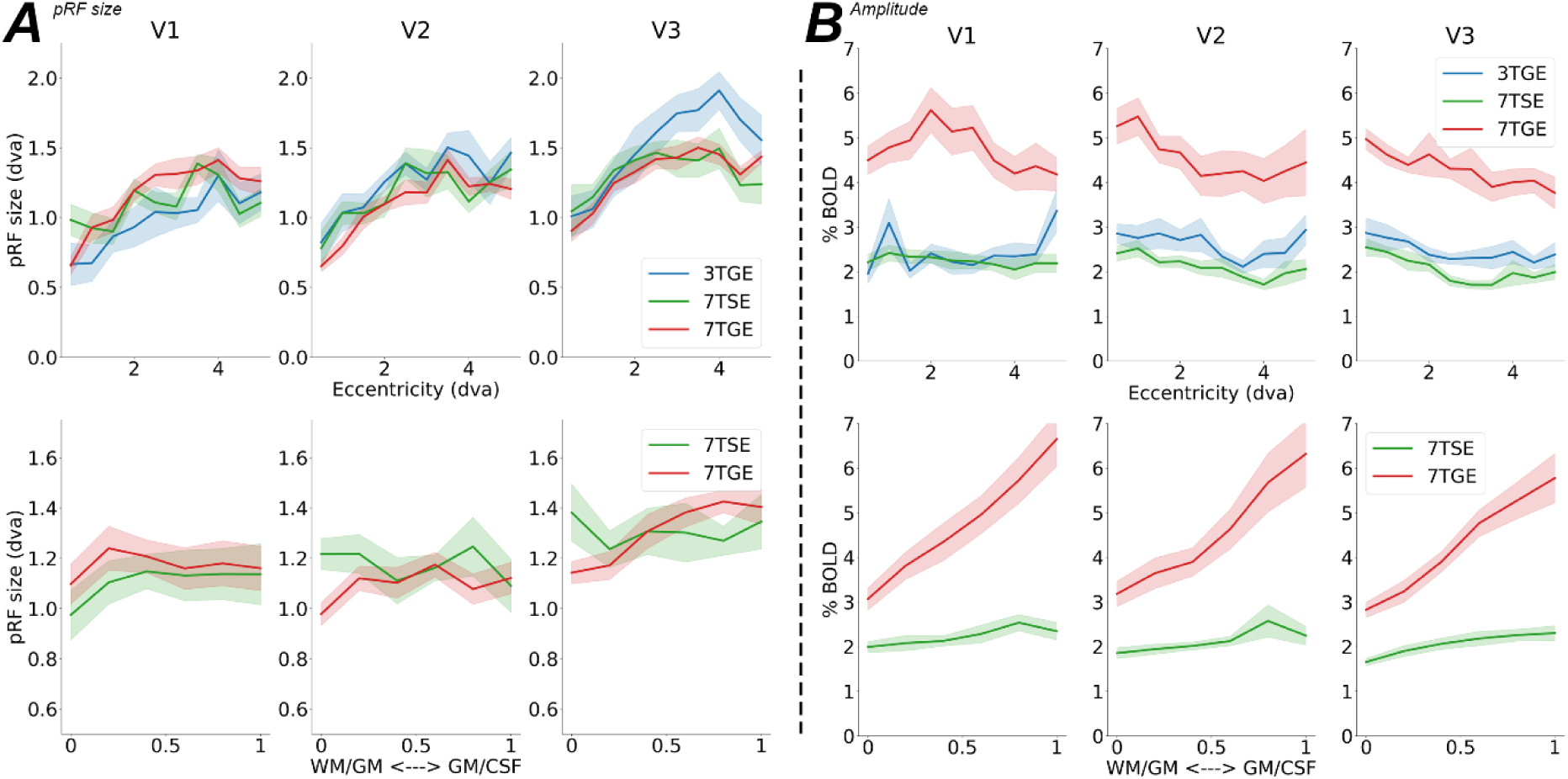
pRF size & amplitude. Mean results of pRF size (A) and BOLD amplitude (B) are shown across eccentricity representations (top panels) and across cortical depth levels (bottom panels) for each MR sequence (colored lines). The shaded area around the line represents the standard error across participants. Abbreviations: dva = degrees visual angle; WM = white matter boundary; GM = grey matter boundary.

**Figure 4.**
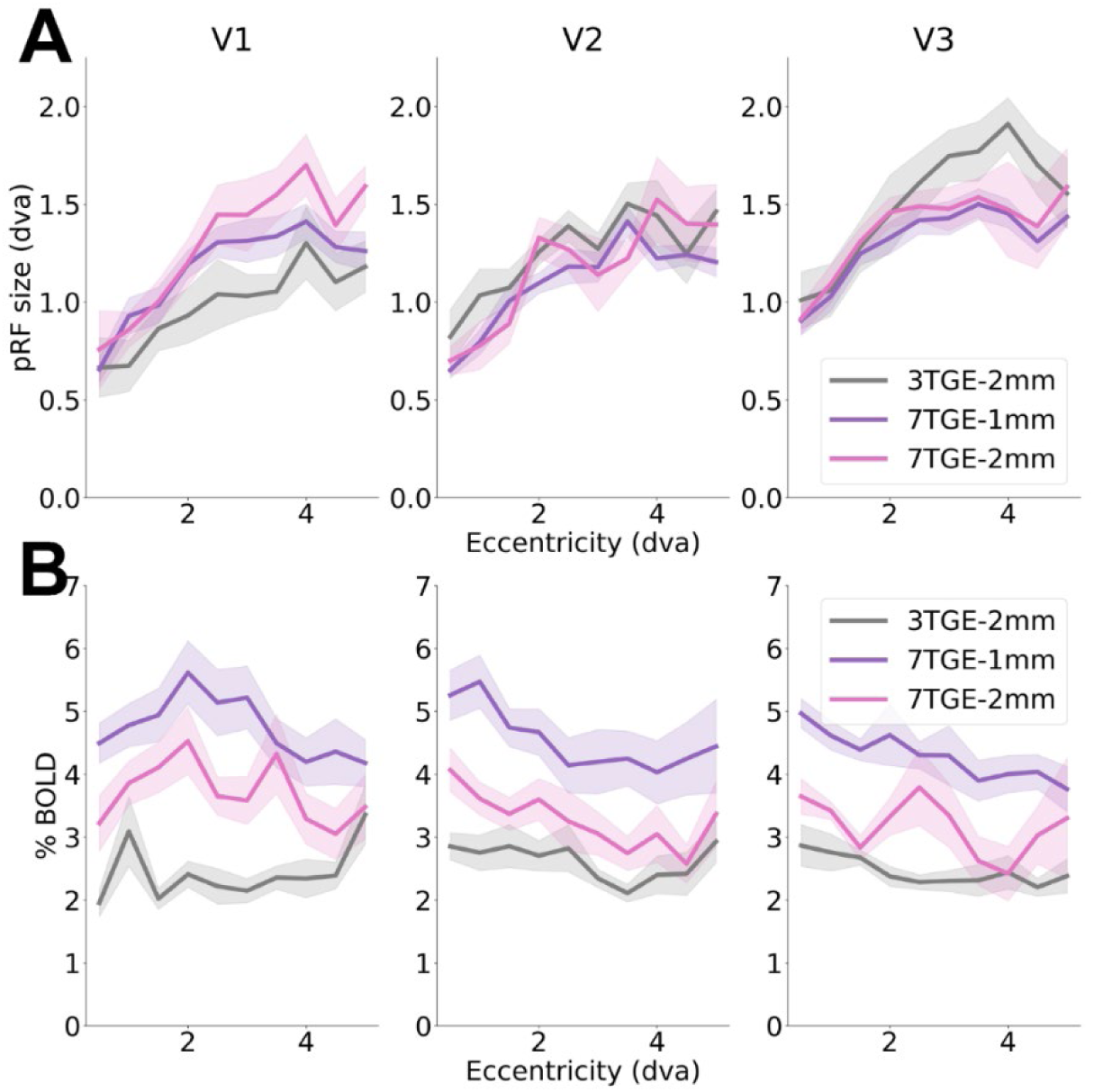
Comparison GE MR sequences. The pRF size (A) and BOLD amplitude (B) estimates are shown for the original (i.e., same as in Figure 2) 7TGE (purple line) and 3TGE MR sequences (grey line). Additionally, the down sampled 2mm isotropic 7TGE MR sequence (pink line) is shown here. Area around the lines represents the standard error across participants.

### PRF size and amplitude

This study aims to reveal vascular contributions to neuronal metrics for fMRI measurements. The previous statistical models presented pRF size dependency on vascular contributions reflected by different MR sequences and spatial voxel categorizations (i.e., visual area, eccentricity representation, and cortical depth level). The current section provides regression analyses on spatial non-specific estimates. First, we performed a linear regression on pRF estimates of different MR sequences. The linear regression was performed with the hypothesized smaller versus larger pRF size MR sequence (e.g., 7TSE versus 7TGE). On average the linear regression slope was *β* = 0.54, s.d. = 0.41, and differed significantly from zero across participants for 7TSE vs 7TGE (t(9) = 6.64, p < 0.001), 7TGE vs 3TGE (t(9) = 4.41, p = 0.004) and 7TSE vs 3TGE (t(9) = 3.33, p = 0.018) (Figure 5A). Additionally, pRF sizes did not correlate with BOLD amplitude *β* = 0.002, s.d. = 0.008 (t(9) = 0.561, p = 0.589). This result indicates that pRF size is not related to the amplitude of the BOLD response (Figure 5b).

**Figure 5.**
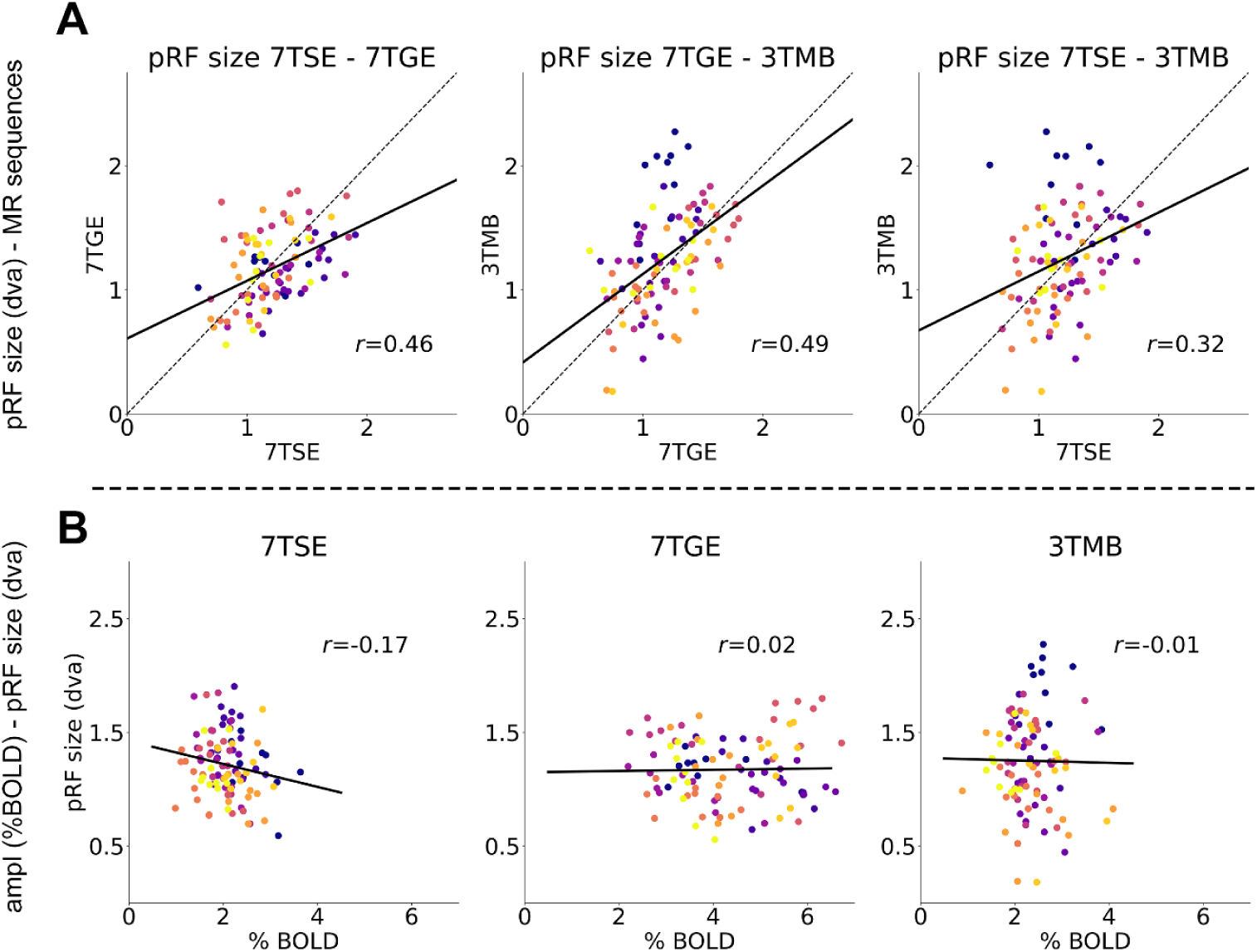
Correlations pRF size and amplitude. A). Scatter plot shows correlations within MR sequences and the fitted linear regression (colored solid line). Perfect correlation is given as a reference by the dashed black line. B) Correlations BOLD amplitude and pRF size within MR sequence (left to right panels). For every panel, each dot is taken as the mean of an eccentricity bin (0-5° visual angle with steps of 0.5° visual angle) for each participant (colors).

## Discussion

### General discussion

Our study investigated the effect of different vascular compartments on neural metrics, such as pRF modeling, for studying brain function using BOLD-fMRI. We utilized T2*-weighted GE and T2-weighted SE sequences at 7T and a multi-band GE sequence at 3T to quantify the contribution of different vascular compartments to pRF estimates within the early visual cortex (V1, V2, and V3). Our findings verify pRF size increases across eccentricity and visual areas, indicating larger receptive fields for peripheral visual field representations and along the cortical hierarchy. Despite expectations, pRF size estimates were not significantly affected by MR sequence or even voxel size, indicating robustness across the different imaging conditions examined here. Consequently, pRF sizes from different MR sequences correlated significantly. The BOLD signal amplitude did not significantly vary across eccentricity representations or visual areas, but was notably affected by the MR sequence with largest signal change seen for the 7TGE MR sequence. Additionally, we observed the well-documented increase in BOLD amplitude from white matter toward the pial surface for the GE MR sequence, but not for the SE MR sequence. Our findings suggest that neural metrics such as the pRF size are not noticeably affected by local vascularization differences as reflected by current scan sequence vessel sensitivities, However, BOLD-fMRI signal amplitude -which is most commonly used to assess brain functioning-appears strongly affected by local vascularization sensitivity.

### Population receptive field sizes

Based on the current understanding of human visual cortex, we expected to find a pRF size increase across eccentricity representation (Harvey and Dumoulin, 2011; Silva et al., 2018), a pRF size increase along the cortical hierarchy from V1 to V3 (Felleman and Van Essen, 1991; Wandell and Winawer, 2015), and a pRF size difference (U-shape) across cortical depth (Fracasso et al., 2016; Self et al., 2019). However, from these hypotheses, only the pRF size increase across eccentricity representations and visual areas could be confirmed. No notable effect was observed in pRF size estimates across cortical depth, which may simply be due to insufficient spatial resolution of the 7TGE (1.0mm isotropic voxel size) and 7TSE (1.5mm isotropic voxel size) MR sequences to resolve a U-shape across cortical depth as compared to previous reports that used a sub-millimeter spatial resolution (Fracasso et al., 2016).However, an amplitude increase was observed across cortical depth for the 7TGE MR sequence, indicative of cortical depth-dependent sensitivity at least to a certain degree. This amplitude increase across cortical depth is caused by an increase in veinous vessel diameter towards the pial surface which determines the GE signal strength greatly. Hence, such amplitude increase across cortical depth was not observed for the 7TSE MR sequence, which is mostly sensitive to micro-vascular compartments. While confirming previous results is important, the main objective of this study was to assess how different vascularization properties, as reflected by varying vessel sensitivities in MR sequences, influence pRF size estimates. Interestingly, all three MR sequences produced comparable pRF size estimates. This aligns with recent findings where no pRF differences were observed between GE BOLD and VASO-CBV at 7 Tesla (Oliveira et al., 2022). In that study, VASO-CBV was used to measure neuronal activity related to microvascular compartments, which are thought to be limited to smaller vessels (Huber et al., 2019, 2017). Consistent with our results, pRF sizes were similar between GE BOLD and VASO-CBV, including the typical increase in pRF size with eccentricity.

The question then arises, if neuronal activity measured by fMRI is influenced by underlying vasculature and specific fMRI sequences have different sensitivities to vascular compartments, why do pRF size estimates remain consistent across MR sequences? We propose several possible explanations. First, pRF size estimates may not depend on local vascularization. PRF calculations are derived from the BOLD signal amplitudes across a set of conditions (e.g., visual field locations), though this relationship is not necessarily linear.

While BOLD signal amplitude varies with vascular compartment size, the relative distribution of amplitudes across conditions—and thus the estimated pRF size—may remain consistent across different vascular compartments. Second, we used a relatively large voxel size for the 7TSE sequence. Our pRF models predicted that voxel size would enlarge pRF sizes and, additionally, predicted that the difference between 7TSE and 7TGE would be marginal due to the larger voxel size of 7TSE compared to 7TGE. The 7TSE voxel size was pre-determined to match the SNR of the 7TGE sequence and may have reduced the likelihood of observing significant differences (Shmuel et al., 2007). Last, the stimulus may have been too small. Current stimulus settings were limited by the projector screen of the 7 Tesla scanner, allowing for a maximal radius of 4.42° visual angle. Possibly, high-resolution mapping stimuli are preferred to assess neuronal population responses in high detail (Prabhakaran et al., 2020). Using relatively small stimuli may lead to blurring of neuronal population responses and a loss in differentiation capacity.

### Neuronal or vascular receptive fields?

The hemodynamic response is influenced by changes in blood flow, volume, and oxygenation levels, all of which are vascular in nature. Vascular factors such as blood vessel density and distribution should have some impact on the spatial extent and organization of pRFs. However, disentangling the relative contributions of neuronal and vascular factors to pRFs is challenging. In the current study, we do not find evidence for a dependency of pRF size on vascular compartment size. Possibly, pRFs primarily reflect the summation of neuronal activity, meaning that they are predominantly neuronal in origin. However, changing the resolution of the 7T gradient echo MR sequence, meaning an eightfold volumetric increase, to match the 3T gradient echo MR sequence did not significantly affect pRF sizes. With volumetric increase, the number of neurons within a voxel naturally increases as well, which could theoretically lead to larger pRF estimates. However, this did not materialize in our findings. Moreover, our predictions were mostly driven by MR-related vascular properties, for which we found no evidence. Hence, it seems that fMRI pRF estimates are somewhat decoupled from the underlying vascular processes. Perhaps, pRF properties produced by the neurovascular coupling behave orthogonally to vascular dynamics, only moderately changing size and shape given the local CBV change. In general, it appears that the relative distribution of signals of a set of conditions (such as coding for a set of visual field locations) is consistent along the vascular tree in early visual cortex. Future studies could address whether this property holds for other types of stimuli and experiments in visual cortex and other brain regions.

## Conclusion

Our study demonstrates that population receptive field (pRF) size estimates in early visual cortex remain consistent across different fMRI sequences, voxel sizes, and magnetic field strengths, despite expectations of vascular compartment type influence. These findings suggest that pRF estimates are primarily neuronal in origin and largely decoupled from the underlying vascular dynamics. While the hemodynamic response is inherently vascular, pRF modeling reflects the neuronal activity more robustly.

## Supplementary Figures

**Supplementary Figure 1.**
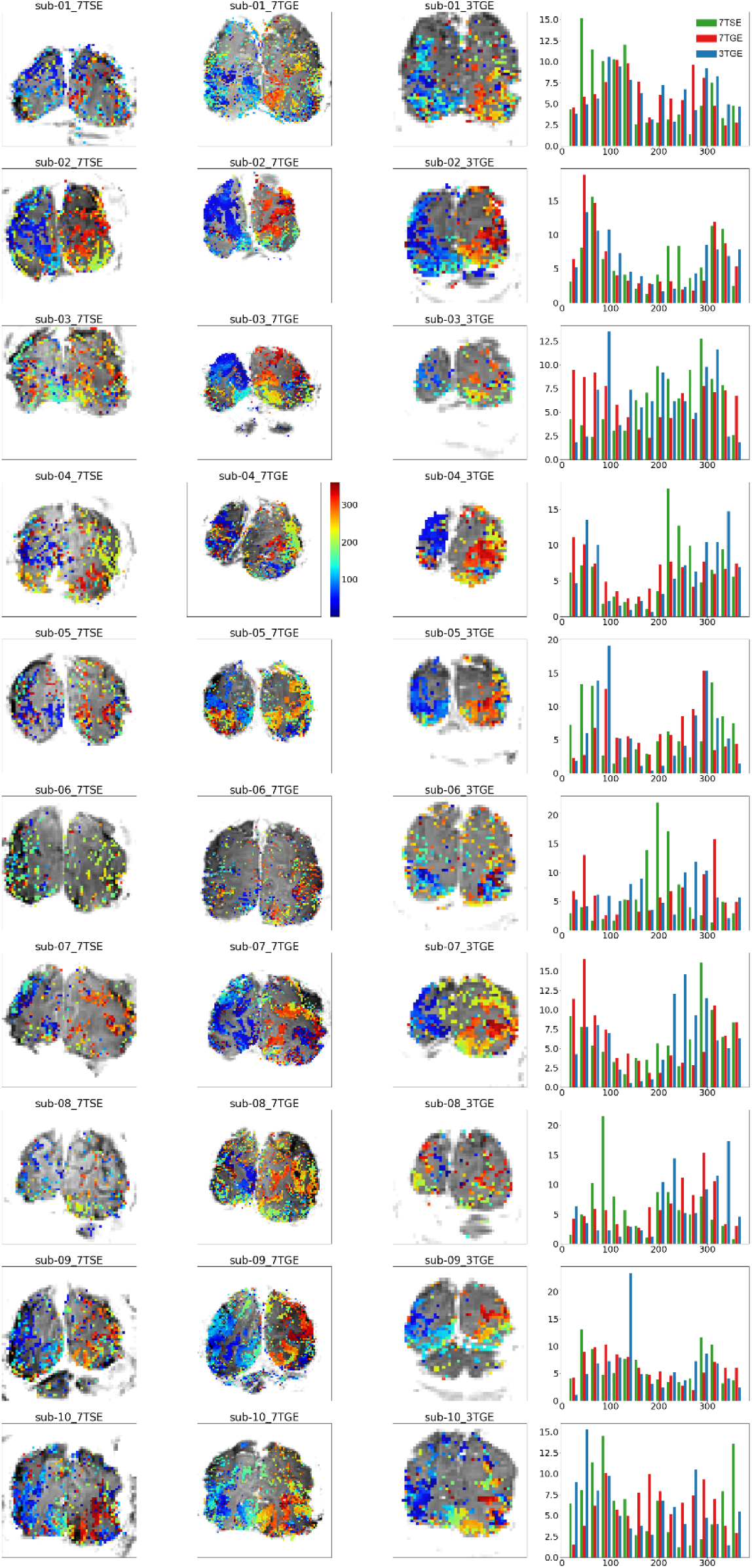
Polar angle results all participants. Polar angle results are shown for each scan sequence (first 3 columns), including a normalized histogram of values (4^th^ column) for all participants (rows).

**Supplementary Figure 2.**
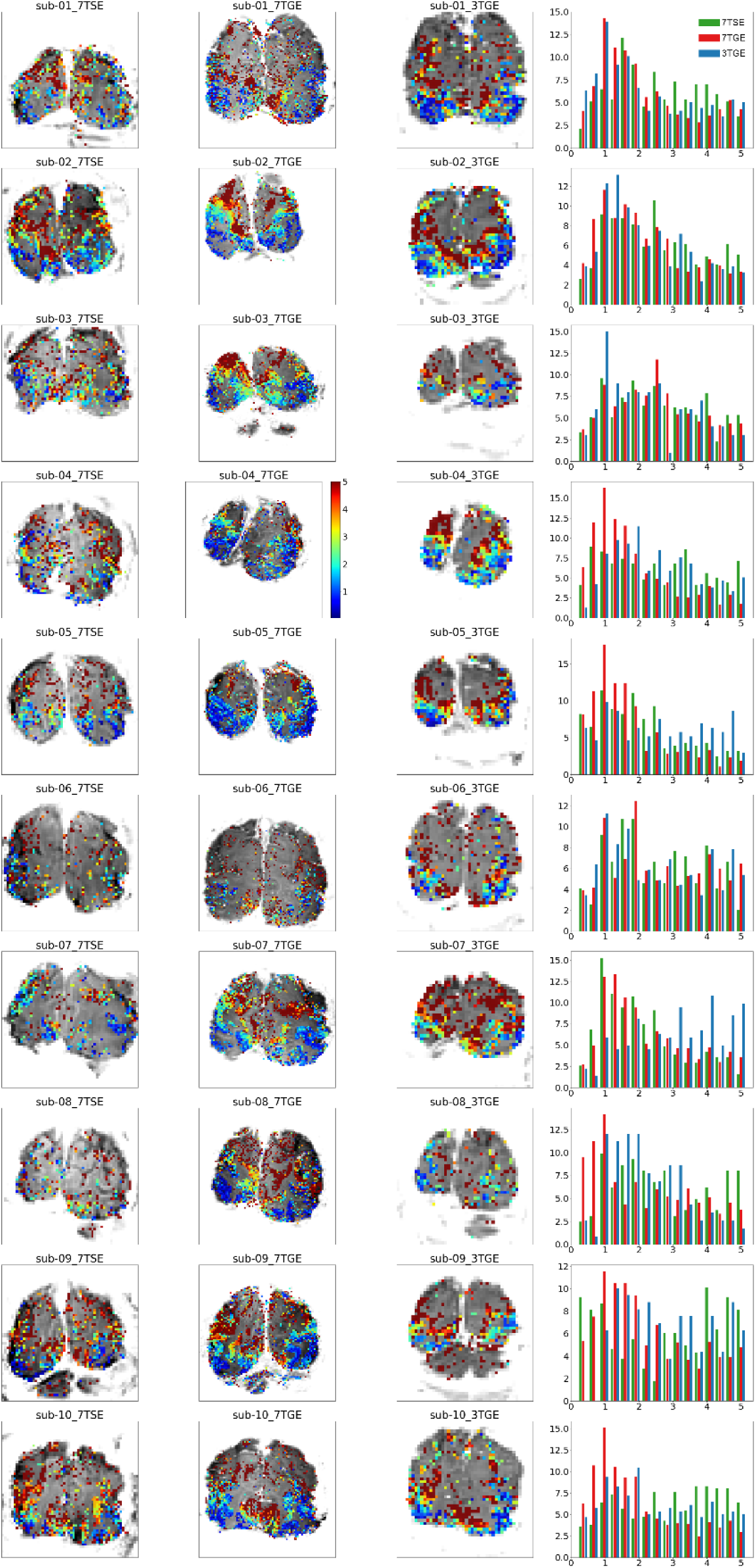
Eccentricity results all participants. Eccentricity results are shown for each scan sequence (first 3 columns), including a normalized histogram of values (4^th^ column) for all participants (rows).

**Supplementary Figure 3.**
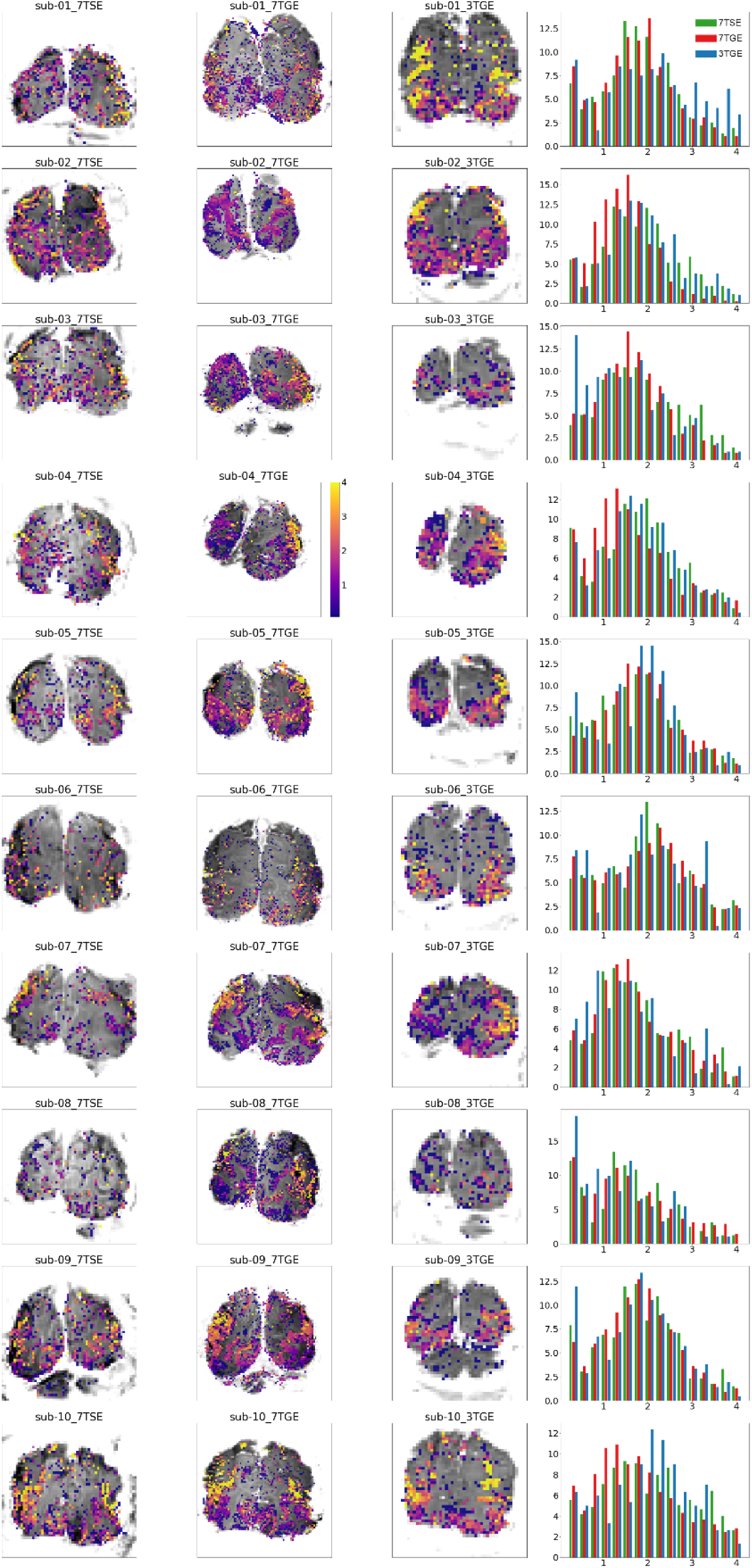
pRF size results all participants. Polar angle results are shown for each scan sequence (first 3 columns), including a normalized histogram of values (4^th^ column) for all participants (rows).

**Supplementary Figure 4.**
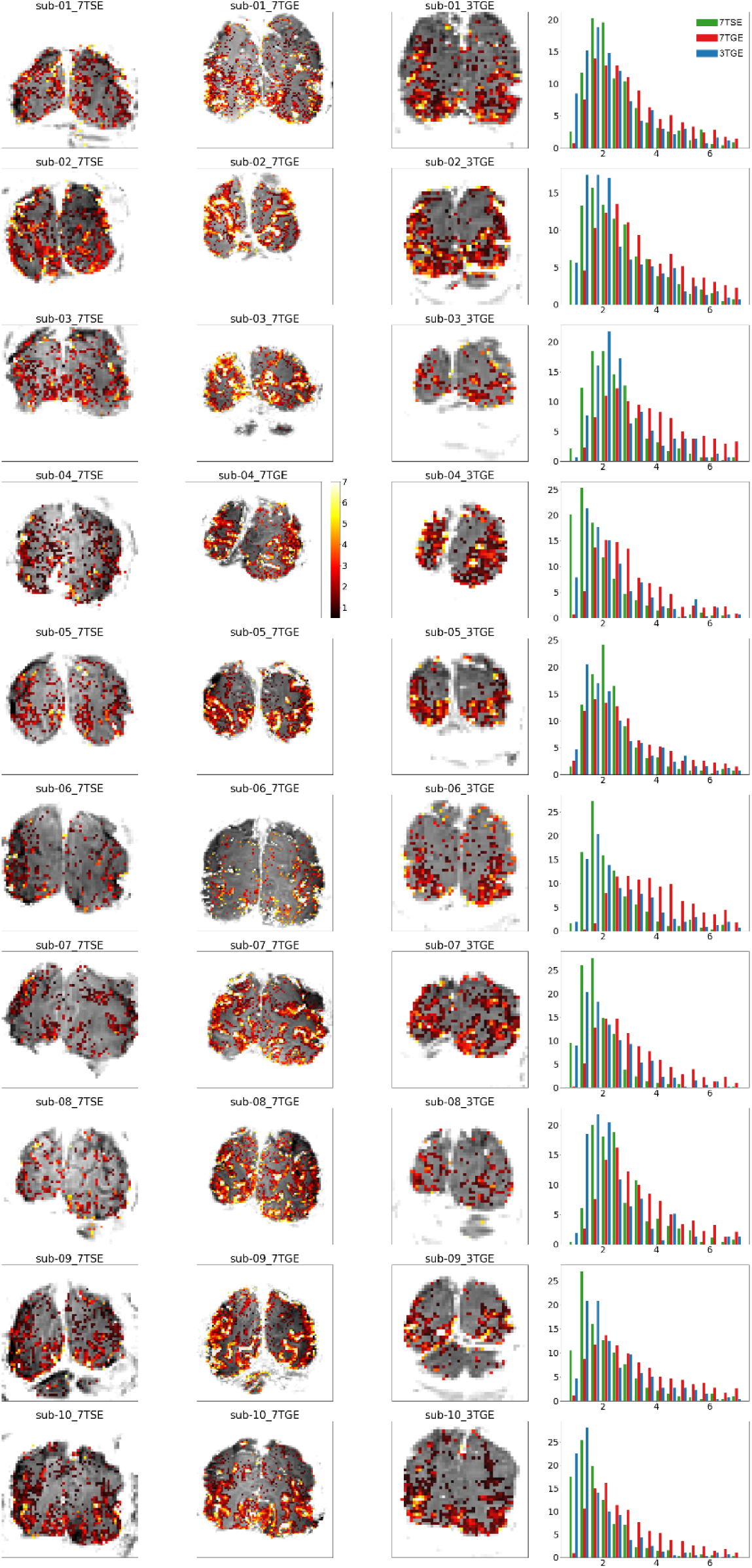
BOLD amplitude results all participants. BOLD amplitude (percent signal change) results are shown for each scan sequence (first 3 columns), including a normalized histogram of values (4^th^ column) for all participants (rows).

**Supplementary Figure 5.**
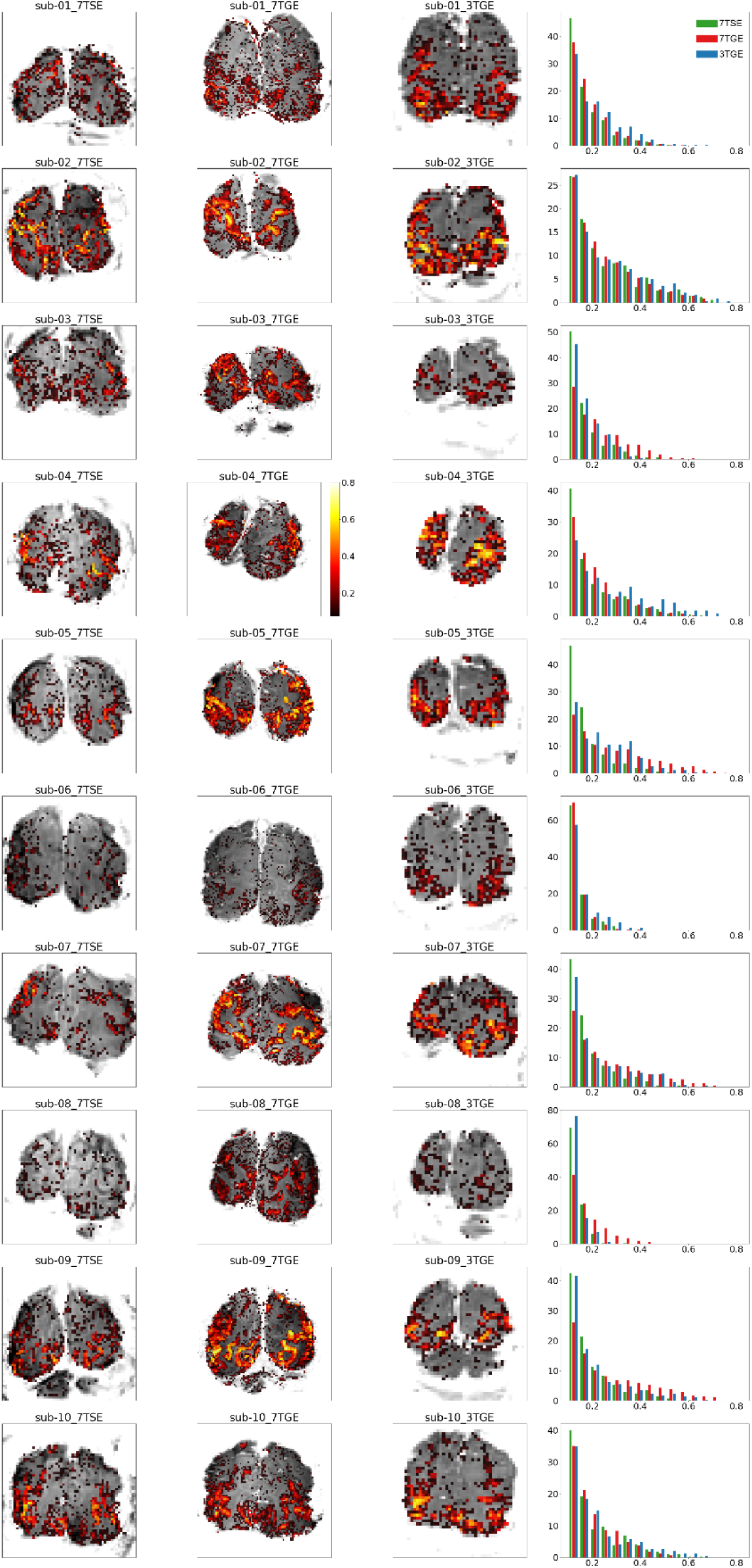
R^2^ pRF fit all participants. R^2^ of the pRF fit is shown for each scan sequence (first 3 columns), including a normalized histogram of values (4^th^ column) for all participants (rows).

